# Cytoplasmic-nuclear incompatibility between wild-isolates of *Caenorhabditis nouraguensis*

**DOI:** 10.1101/083154

**Authors:** Piero Lamelza, Michael Ailion

## Abstract

How species arise is a fundamental question in biology. Species can be defined as populations of interbreeding individuals that are reproductively isolated from other such populations. Therefore, understanding how reproductive barriers evolve between populations is essential for understanding the process of speciation. Hybrid incompatibility (e.g. hybrid sterility and lethality) is a common and strong reproductive barrier in nature, but few studies have molecularly identified its genetic basis. Here we report a lethal incompatibility between two wild-isolates of the nematode *Caenorhabditis nouraguensis.* Hybrid inviability results from the incompatibility between a maternally inherited cytoplasmic factor from each strain and a recessive nuclear locus from the other. We have excluded the possibility that maternally inherited endosymbiotic bacteria cause the incompatibility by treating both strains with tetracycline and show that hybrid death is unaffected. Furthermore, cytoplasmic-nuclear incompatibility commonly occurs between other wild-isolates, indicating that this is a significant reproductive barrier within *C. nouraguensis*. We hypothesize that the maternally inherited cytoplasmic factor is the mitochondrial genome and that mitochondrial dysfunction underlies hybrid death. This system has the potential to shed light on the dynamics of divergent mitochondrial-nuclear coevolution and its role in promoting speciation.

## INTRODUCTION

How species arise is a fundamental and still unanswered question in biology. Under the biological species concept, species consist of populations of interbreeding individuals that are reproductively isolated from other such populations (Mayr 1942). Thus, to understand speciation, we must learn how reproductive barriers evolve between populations. Post-zygotic reproductive barriers are commonly found in nature, and occur when hybrid progeny are relatively unfit in comparison to their parents and serve as inefficient bridges for gene flow between populations. Hybrids can be extrinsically unfit, in that they are maladapted to their environment (e.g. hybrids exhibit an intermediate phenotype which is unfit in parental environments) or intrinsically unfit, in that they are developmentally abnormal (e.g. hybrids are sterile or inviable) (Coyne and Orr 2004).

The Bateson-Dobzhansky-Muller (BDM) model hypothesizes that hybrids are intrinsically unfit due to incompatible gene combinations. In its simplest form, the model predicts that at least two genetic loci, each having evolved independently in one of two divergent lineages, have deleterious epistatic interactions in hybrids. This model has gained support by the molecular identification of genes required for hybrid dysfunction in several genera (Presgraves 2010). Identifying these genes and the natural forces that drive their evolution is one of the major objectives of speciation genetics. Darwin suggested that differential ecological adaptation by natural selection was the major driving force for speciation. Some of the molecularly identified “incompatibility genes” do indeed show signs of selection (Ting 1998; Presgraves *et al.* 2003; Barbash *et al.* 2004; Brideau *et al.* 2006; Oliver *et al.* 2009; Chae *et al.* 2014; Phadnis *et al.* 2015), but do not always have a clear role in promoting ecological adaptation (Tao *et al.* 2001; Ferree and Barbash 2009; Phadnis and Orr 2009; Seidel *et al.* 2011). However, there are currently only a handful of known incompatibility genes from a limited number of genera. Additional studies from a wider range of taxa are needed to gain a better understanding of the evolutionary forces that drive speciation.

Some studies on the genetic basis of hybrid incompatibility have focused on strong post-zygotic reproductive barriers between well-defined species. These studies reveal that many genetic variants contribute to dysfunction of hybrids (Coyne and Orr 1998), but provide little insight into the dynamics of the accumulation of such variants or their relative roles in initiating speciation. For example, theoretical work indicates that the number of genetic incompatibilities increases greater than linearly with the number of genetic differences between two lineages (Orr 1995). Therefore, a small number of genetic incompatibilities may initially reduce gene flow and promote genetic divergence between populations, while others evolve after strong reproductive barriers have already been established. Given this, studies of incomplete post-zygotic barriers between young species or divergent populations within species are essential to understand the evolutionary forces that may initiate speciation.

Despite the paucity of molecularly identified incompatibility genes, the segregation of deleterious phenotypes in a number of interspecific hybridizations indicates that incompatibilities between cytoplasmic and nuclear genomes occur frequently (Ellison and Burton 2008; Ellison *et al.* 2008; Sambatti *et al.* 2008; Arnqvist *et al.* 2010; Ross *et al.* 2011; Aalto *et al.* 2013). Furthermore, several studies have definitively mapped these incompatibility loci to the mitochondrial genome and nuclear genes with mitochondrial functions (Lee *et al.* 2008; Chou *et al.* 2010; Luo *et al.* 2013; Meiklejohn *et al.* 2013; Huang *et al.* 2015). Aerobic eukaryotic organisms rely on mitochondria to generate energy required for diverse biological processes. The mitochondrial genome encodes a small fraction of mitochondrial proteins, but nuclear genes are required for proper replication, transcription, and translation of mtDNA as well as other mitochondrial proteins (Gustafsson *et al.* 2016). Given the interdependence of the nuclear and mitochondrial genomes, they are expected to coevolve by the accumulation of compatible mutations that maintain mitochondrial function. By extension, distinct lineages that undergo unique mitochondrial-nuclear coevolution may be incompatible and result in mitochondrial dysfunction. Several theories have been proposed to explain what drives the rapid coevolution of these two genomes, including adaptation to different carbon sources (Lee *et al.* 2008), arms races between the genomes caused by genetic conflict over the relative fitness of males and females (Fujii *et al.* 2011), and the accumulation of deleterious mtDNA mutations and the evolution of compensatory nuclear variants that rescue mitochondrial function (Rand *et al.* 2004; Oliveira *et al.* 2008; Osada and Akashi 2012). However, given the scarcity of molecularly identified cases of mitochondrial-nuclear incompatibilities, additional studies are required to form more complete theories regarding the forces that drive their evolution.

Here we report incompatibility between the cytoplasmic and nuclear genomes of two distinct wild-isolates of the male-female nematode *Caenorhabditis nouraguensis.* Cytoplasmic-nuclear incompatibility is not specific to these two strains, but is also observed upon hybridization of other distinct wild-isolates of *C. nouraguensis*, indicating that this is a naturally widespread reproductive barrier within the species. The cytoplasmic-nuclear incompatibility we identify between strains of *C. nouraguensis* may provide an excellent opportunity for a detailed study of mitochondrial-nuclear incompatibility, the forces that drive the coevolution of these genomes, and their possible role in speciation.

## MATERIALS AND METHODS

### Strain isolation and maintenance

All strains of *C. nouraguensis* used in this study were derived from single gravid females isolated in 2009 or 2011 from rotten fruit or flowers found in French Guiana (Kiontke *et al.* 2011), and have not been subjected to further inbreeding. Strains were kindly provided by Marie-Anne Felix (“JU” prefix) and Christian Braendle (“NIC” prefix). Strain stocks were stored at −80°C. Thawed strains were maintained at 25°C on standard NGM plates spread with a thin lawn of OP50 bacteria (Brenner 1974).

### Hybridizing JU1825 and NIC59

To quantify inviability, we crossed one virgin L4 female and male, with 10-15 replicates for each cross. The edge of each plate was coated with a palmitic acid solution (10 mg/mL in 95% ethanol) and allowed to air dry, resulting in a physical barrier that helps prevent worms from leaving the plate’s surface. The plates were placed at 25°C overnight, during which the worms matured to adulthood and began mating. The next day, each female-male couple was placed onto a new plate streaked with OP50 and rimmed with palmitic acid. Each couple was then allowed to mate and lay eggs for 5 hours at 25°C, and then were permanently removed. The embryos laid within those 5 hours were counted immediately. Approximately 17 hours later, we counted the number of embryos that failed to hatch per plate. These unhatched embryos were scored as dead since *C. nouraguensis* embryogenesis is normally completed within 13 hours at 25°C (data not shown). We defined the percentage of embryonic lethality as the number of unhatched embryos divided by the total number of embryos laid. Approximately 20 hours later, we placed the plates at 4°C for an hour and then counted the number of healthy L4 larvae and young adults per plate. We defined the percentage of viable progeny as the total number of L4 larvae and young adults divided by the total number of embryos laid.

### Determining cytoplasmic-nuclear compatibility between various strains of C. nouraguensis

To test for an incompatibility between one strain’s cytoplasm and another strain’s nuclear genome, we compared the viabilities of backcrosses that differ only in the F1 hybrid female’s cytoplasmic genotype (e.g. (NIC59); NIC59/JU1837 F1 female × JU1837 male vs (JU1837); NIC59/JU1837 F1 female × JU1837 male, Figure 3A). The genotype is designated by the following nomenclature: (cytoplasmic genotype); nuclear genotype. The cytoplasmic genotype indicates genetic elements that are only maternally inherited, such as the mitochondrial genome. We performed a Fisher’s exact test to determine whether there were significant differences in the proportions of viable and inviable F2 progeny between the two types of crosses. We also calculated the relative viability of the two crosses (e.g. the percent viability of the (NIC59); NIC59/JU1837 F1 female × JU1837 male cross divided by the percent viability of (JU1837); NIC59/JU1837 F1 female × JU1837 male cross). Cytoplasmic-nuclear combinations that show a statistically significant difference in viabilities between the two types of crosses and a relative viability <1 were considered to be cytoplasmic-nuclear incompatibilities. Three biological replicates were performed for each cytoplasmic-nuclear combination except for JU1825 cytoplasmic - NIC24 nuclear and JU1825 cytoplasmic - NIC54 nuclear, which have four replicates each. For each biological replicate, 10 F1 hybrid L4 females were crossed to 10 L4 males on the same plate overnight at 25°C. The next day, they were moved to a new plate and allowed to lay embryos at 25°C for 1 hour. The parents were then removed and the percent viable progeny and embryonic lethality were calculated as described in the previous section of the Materials and Methods. The heat map used to visualize the median relative viability for each cytoplasmic nuclear combination was made using the heatmap.2 function from the gplot package in R.

### Molecular Methods

To determine if either JU1825 or NIC59 are infected with *Wolbachia*, we performed PCR on crude lysates of both strains using degenerate primers targeted against two genes that are conserved in *Wolbachia* (Baldo *et al.* 2006). Specifically, we attempted to detect *gatB* (gatB_F1 with M13 adapter, TGTAAAACGACGGCCAGTGAKTTAAAYCGYGCAGGBGTT, and gatB_R1 with M13 adapter, CAGGAAACAGCTATGACCTGGYAAYTCRGGYAAAGATGA) and *fbpA* (fbpA_F3, GTTAACCCTGATGCYYAYGAYCC, and fbpA_R3, TCTACTTCCTTYGAYTCDCCRCC). As controls, we performed PCR on squash preps of *Drosophila melanogaster*w^1118^ mutant strains (Bloomington stock number 3605) that were infected or not infected with *Wolbachia. Drosophila melanogaster* strains were kindly provided by the laboratories of Harmit Malik and Leo Pallanck.

### Tetracycline treatment of JU1825 and NIC59

Both JU1825 and NIC59 were passaged on 50 ug/mL tetracycline NGM plates streaked with OP50 for nine generations. Both strains were treated by placing 10 L4 females and 10 L4 males on a fresh tetracycline plate and allowing them to mate and produce the next generation of L4 progeny, which were then moved to a fresh tetracycline plate. Tetracycline plates were made by allowing NGM plates with OP50 lawns to soak up a mixture of tetracycline and 1x M9. The plates were left uncovered at room temperature until dry, and then used the following day.

### Statistics

P values were determined using R (v 3.2.5). Several statistical tests were used (Kruskal-Wallis test followed by Dunn’s test, and Fisher’s exact test). When we performed several comparisons on the same dataset, we used the Bonferroni method to correct p-values for multiple testing. Most plots were made using the ggplot2 package in R.

### Data Availability

The authors state that all data necessary for confirming the conclusions presented in the article are represented fully within the article and Supplemental Material.

## RESULTS

### Two strains of *C. nouraguensis* exhibit F2 hybrid breakdown

Two strains of *C. nouraguensis*, JU1825 and NIC59, were derived from single gravid females that were isolated approximately 112 kilometers apart in French Guiana (Kiontke *et al.* 2011). Both of these strains were designated as belonging to *C. nouraguensis* based on having highly similar ITS2 rDNA sequences, which serves as a good species barcode within the *Caenorhabditis* genus, and because they produced many viable F1 offspring when crossed (Kiontke *et al.* 2011; Félix *et al.* 2014). We found that both strains produce high numbers of viable progeny in intra-strain crosses. We have also confirmed the previous finding of F1 hybrid viability by crossing NIC59 females to JU1825 males, and vice versa, showing that the F1 hybrids resulting from these inter-strain crosses exhibit levels of viability comparable to those seen in intra-strain crosses (Figure 1).

**Figure 1.**
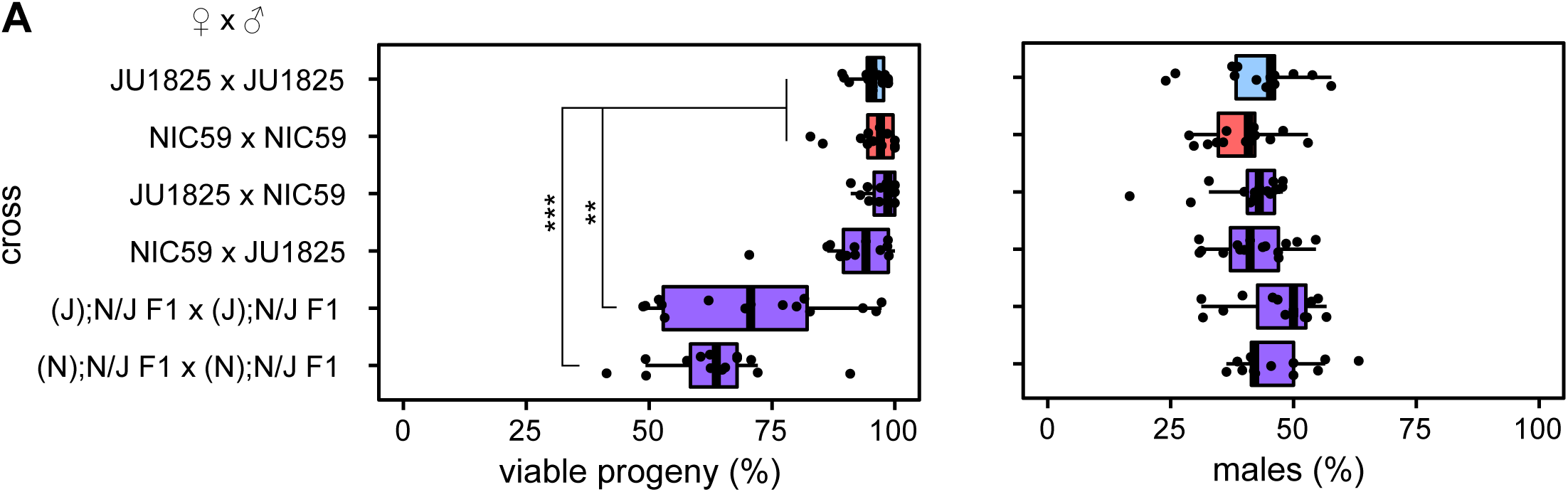
JU1825 and NIC59 exhibit F2 hybrid breakdown. Crosses are listed on the y-axis. Letters in parentheses to the left of a semi-colon denote the cytoplasmic genotype of an individual (e.g. “(J)” individuals have a JU1825 cytoplasmic genotype), while letters to the right of a semicolon denote the genotypes of all autosomal loci (i.e. “N/J” individuals are heterozygous NIC59/JU1825 throughout the autosomes). Only (J); N/J F1 × (J); N/J F1 and (N); N/J F1 × (N); N/J F1 crosses exhibit a significant decrease in the percentage of viable progeny (P<0.01 and P<0.001, respectively). There are no significant differences in the percentages of viable males between crosses (P>0.05). N=14 or 15 plates per cross. All p-values were calculated by a Kruskal-Wallis test followed by Dunn’s test.

However, not all reproductive barriers act in the F1 generation. There are many cases of F2 hybrid breakdown, in which reduction of hybrid fitness begins to manifest itself in the F2 generation due to recessive incompatibility loci (Masly *et al.* 2006; Bikard *et al.* 2009; Stelkens *et al.* 2015). To test for F2 hybrid inviability, we mated hybrid F1 siblings derived from either JU1825 female × NIC59 male crosses, or from NIC59 female × JU1825 male crosses, and assayed the F2 generation for reductions in fitness. These F1 hybrids are referred to as “(J); N/J” and “(N); N/J” respectively, where the genotype is designated by the following nomenclature: (cytoplasmic genotype); nuclear genotype. The cytoplasmic genotype indicates genetic elements that are only maternally inherited, such as the mitochondrial genome. We found that both types of F1 sibling crosses resulted in a significant decrease in the percentage of viable progeny, with on average only 71% and 63% of F2 embryos maturing to the L4 or young adult stage (Figure 1). These results indicate that there are divergent genomic loci between NIC59 and JU1825 that cause inviability only when they become homozygous in F2 hybrids. Additionally, there is no difference in sex-specific mortality in hybrids in comparison to intra-strain crosses which implies that these loci are autosomally linked, as we show later (Figure 1).

### Incompatibilities between cytoplasmic and nuclear genomes cause F2 inviability

To further understand the genetic architecture of hybrid breakdown between JU1825 and NIC59, we tested whether maternally or paternally inherited factors are required for F2 inviability. We reasoned that backcrossing F1 females to parental males would test whether maternal factors are required for reduced hybrid fitness, while backcrossing F1 males to parental females would test whether paternal factors are required. For example, backcrossing F1 hybrid females to JU1825 males will result in an F2 population with a 50% chance of being heterozygous (NIC59/JU1825) and a 50% chance of being homozygous (JU1825/JU1825) for any given autosomal locus. Therefore, this cross will test for maternally deposited NIC59 factors that are incompatible with homozygous JU1825 autosomal loci. The same logic can be applied to crosses of F1 hybrid males to parental strain females.

All instances of backcrossing F1 hybrid males to parental strain females resulted in levels of F2 viability similar to those observed in parental strains. Therefore, paternal factors do not have a major effect on F2 inviability (Figure 2A). Only two crosses consistently resulted in significantly reduced viability. The first is when (N); N/J F1 females were crossed to JU1825 males, with on average only 36% of F2 hybrids maturing to the L4 or young adult stage. This cross implies that there are maternally derived NIC59 factors distributed to F2 embryos, and these factors are incompatible with recessive JU1825 nuclear loci. The second is when (J); N/J F1 females are crossed to NIC59 males, with on average only 52% of the F2 hybrids maturing to the L4 or young adult stage (Figure 2B). This cross implies that there are also maternally derived JU1825 factors distributed to F2 embryos, and these factors are incompatible with recessive NIC59 nuclear loci. The viability of (J); N/J F1 female × JU1825 male crosses can also be significantly reduced in comparison to intra-strain crosses, but varies within and between experiments (Figure S1).

**Figure 2.**
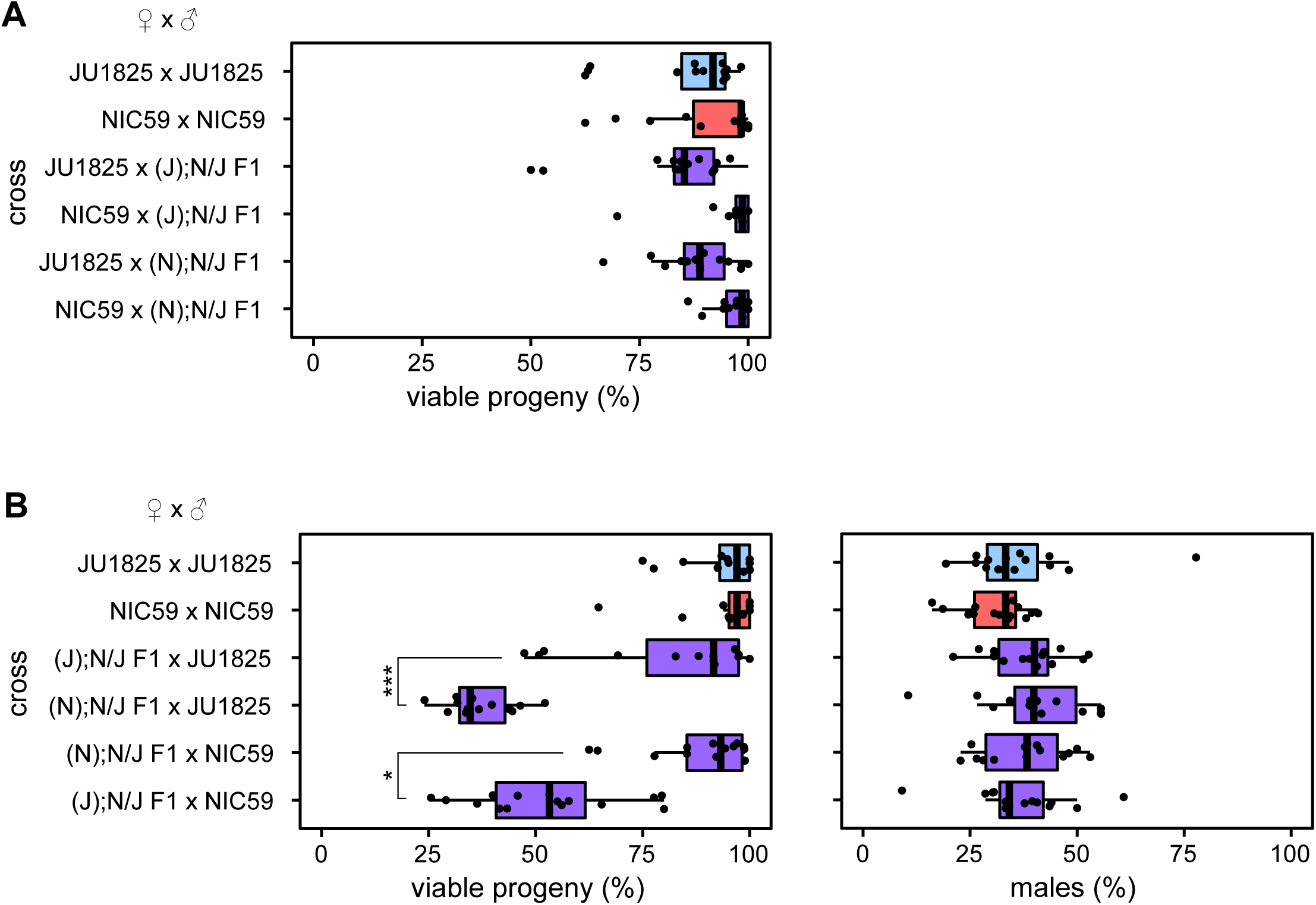
F2 inviability involves a maternal cytoplasmic effect. **(A)** There is no significant difference in the percentage of viable progeny between any of the F1 hybrid male backcrosses and intra-strain crosses (P>0.05). **(B)** Backcrossing hybrid females to parental strain males reveals that only (N); N/J F1 female × JU1825 male crosses and (J); N/J F1 female × NIC59 male crosses exhibit a significant decrease in the percentage of viable progeny in comparison to intrastrain crosses (P<0.001). (N); N/J F1 female × JU1825 male crosses have significantly decreased viability in comparison to (J); N/J F1 female × JU1825 male crosses (P<0.001). Additionally, (J); N/J F1 female × NIC59 male crosses consistently have significantly decreased viability in comparison to (N); N/J F1 female × NIC59 male crosses (P<0.05). The viability of (J); N/J F1 female × JU1825 males can differ significantly between experiments (Figure S1). There are no significant differences in the proportion of viable males between the crosses (P>0.05). N=14 or 15 plates per cross. All p-values were calculated by a Kruskal-Wallis test followed by Dunn’s test.

Interestingly, the F1 female backcross experiments show that almost identical crosses, which differ only in the cytoplasmic genotype of the F1 female, have significantly different rates of F2 viability. For instance, (N); N/J F1 female × JU1825 male crosses consistently have significantly lower F2 viability than (J); N/J F1 female × JU1825 male crosses (Figure 2, Figure S1). Similarly, (J); N/J F1 female × NIC59 male crosses consistently have significantly lower F2 viability than (N); N/J F1 female × NIC59 male crosses (Figure 2). These F1 hybrid females are expected to be genotypically identical at all nuclear loci, suggesting that something other than the F1 nuclear genome encodes maternal factors that lead to F2 inviability. One possible model to explain the differences in these backcrosses is that the mitochondrial genome is the maternally inherited factor that is incompatible with recessive nuclear loci in the F2 generation. For example, all F2 progeny from (N); N/J F1 female × JU1825 male crosses will inherit only NIC59 mtDNA, which may be incompatible with nuclear loci homozygous (or hemizygous) for JU1825 alleles resulting in inviability (Figure 6A). In comparison, all F2 progeny from (J); N/J F1 female × JU1825 male crosses will inherit only JU1825 mtDNA, which should be compatible with the JU1825 nuclear genome and therefore not result in the same inviability. The same logic can be applied to the (J); N/J F1 female × NIC59 male and (N); N/J F1 female × NIC59 male crosses. Therefore, we hypothesize that F2 inviability is the result of two distinct cytoplasmic-nuclear incompatibilities, one between the NIC59 mitochondrial genome and recessive JU1825 nuclear loci, and another between the JU1825 mitochondrial genome and recessive NIC59 nuclear loci.

### The nuclear incompatibility loci are linked to autosomes

Nematodes commonly have an XX (female) and XO (male) sex determining mechanism (Pires-daSilva 2007). The F1 hybrid female backcross experiments reveal that there is no difference in sex-specific mortality in hybrids in comparison to intra-strain crosses (Figure 2B). However, given the expected genotypes of their F2 populations, these backcrosses on their own do not allow us to determine whether the nuclear incompatibility loci are autosomally or X-linked. In the previous section, we concluded that the inviability of the F2 progeny derived from (N); N/J F1 female × JU1825 male crosses is the result of a genetic incompatibility between the NIC59 mitochondrial genome and nuclear loci homozygous (or hemizygous) for JU1825 alleles. If this is true, it is reasonable to assume that the same genetic incompatibility occurs in (N); N/J F1 female × (N); N/J F1 male crosses (Figure 1). In this F1 sibling cross, if the JU1825 nuclear incompatibility locus were autosomally linked, both sexes would suffer equal rates of inviability. However, if the nuclear incompatibility locus were linked to the X-chromosome, then we would expect a significant decrease in the proportion of viable males in comparison to intra-strain crosses (Figure S2). However, we observe no significant difference in the proportion of viable males for the (N); N/J F1 female × (N); N/J F1 male cross (Figure 1). Therefore, given the data from the F1 female backcrosses and the F1 sibling crosses, we conclude that the JU1825 nuclear incompatibility locus is autosomally linked. A similar line of reasoning indicates that the NIC59 nuclear incompatibility locus is also autosomally linked.

### Endosymbiotic bacteria do not cause hybrid inviability

We hypothesize that mitochondrial genomes are responsible for the cytoplasmic component of the hybrid incompatibility between NIC59 and JU1825. However, we also considered whether endosymbiotic bacteria of the *Rickettsiales* order could be involved. Within this order, bacteria of the *Wolbachia* genus are known to infect certain species of nematodes, and are transmitted to host progeny through female gametes (Werren *et al.* 2008). Furthermore, hybrid lethality in inter-strain and interspecies crosses is sometimes caused by infection with divergent *Wolbachia* strains (Bourtzis *et al.* 1996; Bordenstein *et al.* 2001). However, we failed to detect conserved genes typically used to genotype diverse strains of *Wolbachia* in either JU1825 or NIC59 using PCR with degenerate primers (Figure S3A). Additionally, treatment of both strains with tetracycline for nine generations failed to rescue hybrid inviability (Figure S3B). Endosymbiotic bacteria within the *Rickettsiales* order are typically susceptible to tetracycline (McOrist 2000; Darby *et al.* 2015). Thus, endosymbiotic bacteria are unlikely to cause the reproductive barrier between NIC59 and JU1825.

### Cytoplasmic-nuclear incompatibility is common within *C. nouraguensis*

We hybridized additional wild-isolates to determine whether cytoplasmic-nuclear incompatibilities represent a common reproductive barrier within *C. nouraguensis*, or whether they are an unusual phenotype only observed in hybridizations between NIC59 and JU1825. Specifically, we tested the compatibility of four cytoplasmic genotypes with seven nuclear genotypes (Figure 3). To test for an incompatibility between one strain’s cytoplasm and another strain’s nuclear genome, we again compared the viabilities of backcrosses that differ only in the F1 hybrid female’s cytoplasmic genotype (Figure 3A). Specifically, we compared the viability of the backcross that combines heterotypic cytoplasmic and nuclear genotypes to the viability of the backcross that combines homotypic cytoplasmic and nuclear genotypes. We calculated the relative viability of the two crosses (i.e. heterotypic combination / homotypic combination), and tested for statistically significant differences (see Materials and Methods). Using the same logic as for our JU1825 × NIC59 crosses, we reasoned that lower viability of the heterotypic cytoplasmic-nuclear combination in comparison to the homotypic cytoplasmic-nuclear combination indicates a cytoplasmic-nuclear incompatibility. Three or four biological replicates were performed for each cytoplasmic-nuclear combination.

**Figure 3.**
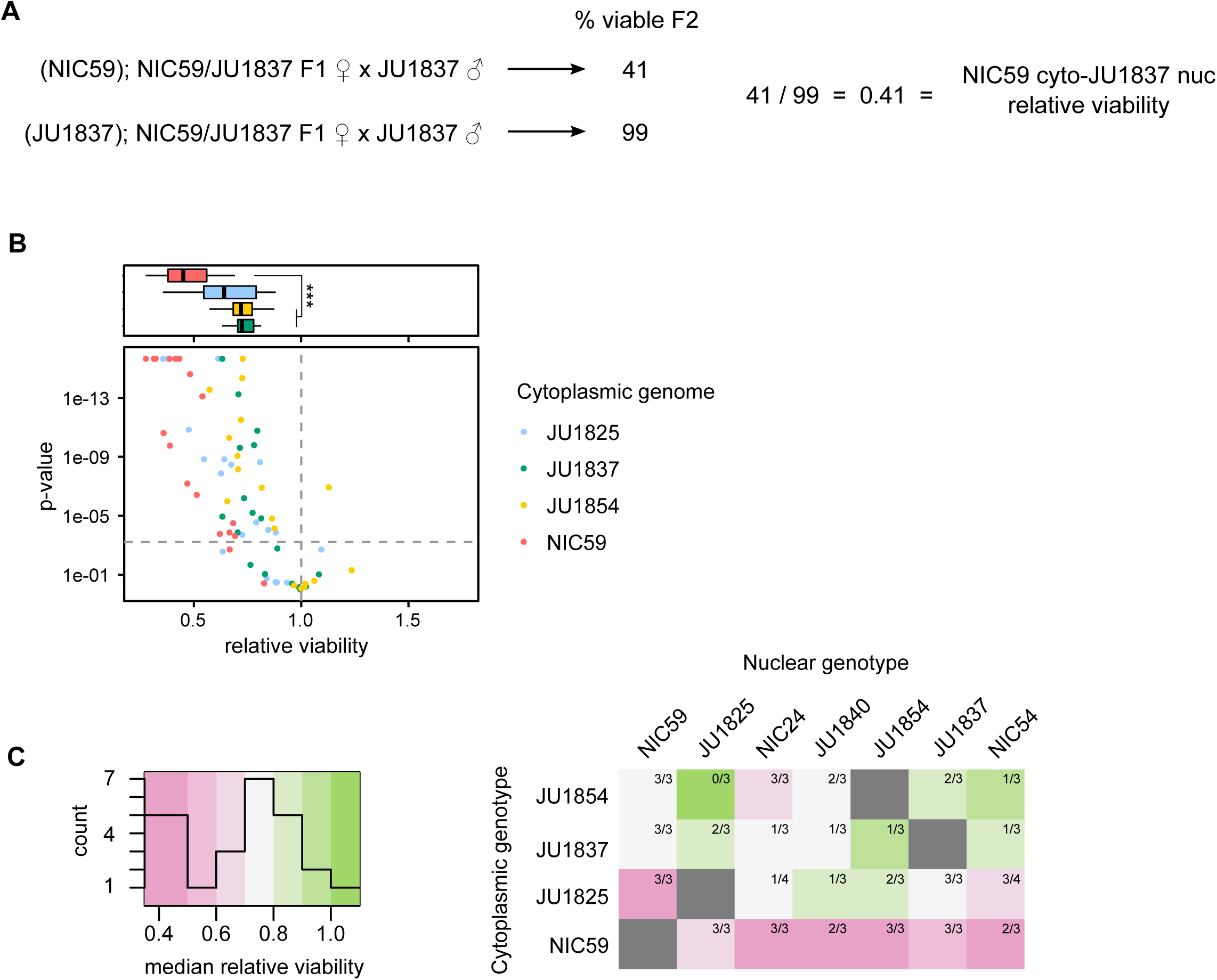
Cytoplasmic nuclear incompatibility is widespread within *C. nouraguensis.* **(A)** To determine whether a particular cytoplasmic-nuclear combination is incompatible, we tested for statistical differences in viability between the F1 female backcross that combines heterotypic cytoplasmic and nuclear genotypes (top cross) and the backcross that combines homotypic cytoplasmic and nuclear genotypes (bottom cross, see Materials and Methods). We also calculated the relative viability of the first cross to the second. **(B)** A scatter plot depicting all the cytoplasmic-nuclear compatibility tests performed. Each point corresponds to a single replicate of a certain cytoplasmic-nuclear combination. Points above the horizontal dashed gray line indicate statistically significant differences in viability between the two types of crosses mentioned in (A) (P<0.0006 after Bonferroni correction, Fisher’s exact test). Points above the horizontal dashed gray line that have a relative viability <1 are considered statistically significant cytoplasmic-nuclear incompatibilities. The color of a point corresponds to the cytoplasmic genotype being tested. All cytoplasmic genotypes tested show an incompatibility with one or more heterotypic nuclear genotypes. See Figure S4 for separate graphs of all combinations. Above the scatterplot are boxplots depicting the relative viabilities of statistically significant cytoplasmic-nuclear incompatibilities. The color corresponds to cytoplasmic genotype tested. Incompatibilities involving the NIC59 cytoplasmic genotype have reduced viability compared to those involving the JU1837 and JU1854 cytoplasmic genotypes (P<0.001, Kruskal-Wallis test followed by Dunn’s test). **(C)** A heatmap depicting the median relative viability for each cytoplasmic-nuclear combination. Each cytoplasmic-nuclear combination shows the proportion of replicates that exhibit significant incompatibilities (e.g. 3 out of 3 replicates exhibit significant incompatibilities for the NIC59 cytoplasm - JU1854 nuclear combination, while only 1 out of 3 replicates exhibit significant incompatibilities for the JU1837 cytoplasm - JU1854 nuclear combination). Each cytoplasmic genotype is consistently incompatible with at least one heterotypic nuclear genotype. The NIC59 cytoplasm has a distinct response to hybridization than the others tested.

Of the 74 cytoplasmic-nuclear tests performed, 50 (67%) exhibited significant incompatibilities (Figure 3B). Additionally, each cytoplasmic genotype was consistently incompatible with at least one heterotypic nuclear genotype (i.e. all replicates for a particular cytoplasmic-nuclear combination indicate a significant incompatibility). However, there are a number of cytoplasmic-nuclear combinations whose replicates are inconsistent with one another (e.g. some replicates indicate a significant incompatibility while others do not) (Figure 3C and Figure S4). This may indicate that the genetic loci required for hybrid inviability are not fixed between the strains, but rather are polymorphisms segregating within each strain. This is consistent with the fact that none of these strains have been formally inbred. Regardless, given their common occurrence in hybridizations between strains of *C. nouraguensis*, we hypothesize that cytoplasmic-nuclear incompatibilities are a significant reproductive barrier within the species.

We generated a heat map to help visualize the median relative viability for each cytoplasmic-nuclear combination (Figure 3C). Strikingly, the NIC59 cytoplasmic genotype exhibits a distinct response to hybridization, being strongly incompatible (i.e. having a low median relative viability) with all of the nuclear genotypes tested. By comparison, the other cytoplasmic genotypes can be relatively compatible with some heterotypic nuclear genotypes or exhibit incompatibilities that are typically weaker than those involving the NIC59 cytoplasmic genotype. Specifically, incompatibilities involving the JU1837 or JU1854 cytoplasmic genotypes have significantly higher relative viability (median=0.72 and 0.71, respectively) in comparison to incompatibilities with the NIC59 cytoplasmic genotype (median=0.45) (Figure 3B). Incompatibilities involving the JU1825 cytoplasm exhibit an intermediate level of relative viability (median=0.64) that is statistically indistinguishable from the other cytoplasmic genotypes (P=0.057 in comparison to NIC59; P=1.0 in comparison to both JU1837 and JU1854). We conclude that the NIC59 cytoplasmic genotype is distinct in terms of the nuclear genotypes it is incompatible with and how severe those incompatibilities are.

### A single BDM incompatibility between a NIC59 cytoplasmic locus and a JU1825 nuclear locus causes embryonic lethality

As previously discussed, the backcross that combines the NIC59 cytoplasmic genotype with JU1825 nuclear genotype (i.e. (N); N/J F1 female × JU1825 male, Figure 2B) results in only ~36% of F2 offspring maturing to the L4 or young adult stage. A more detailed characterization of F2 inviability shows that ~50% of F2 offspring fail to complete embryogenesis (Figure 4A). Of the remaining half that complete embryogenesis, ~33% fail to mature to the L4 or young adult stage (data not shown). In comparison, (J); N/J F1 female × JU1825 male crosses result in low levels of embryonic lethality, similar to parental crosses. These data are consistent with F2 embryonic lethality resulting from a single BDM incompatibility between a NIC59 cytoplasmic locus and a single homozygous JU1825 autosomal locus.

**Figure 4.**
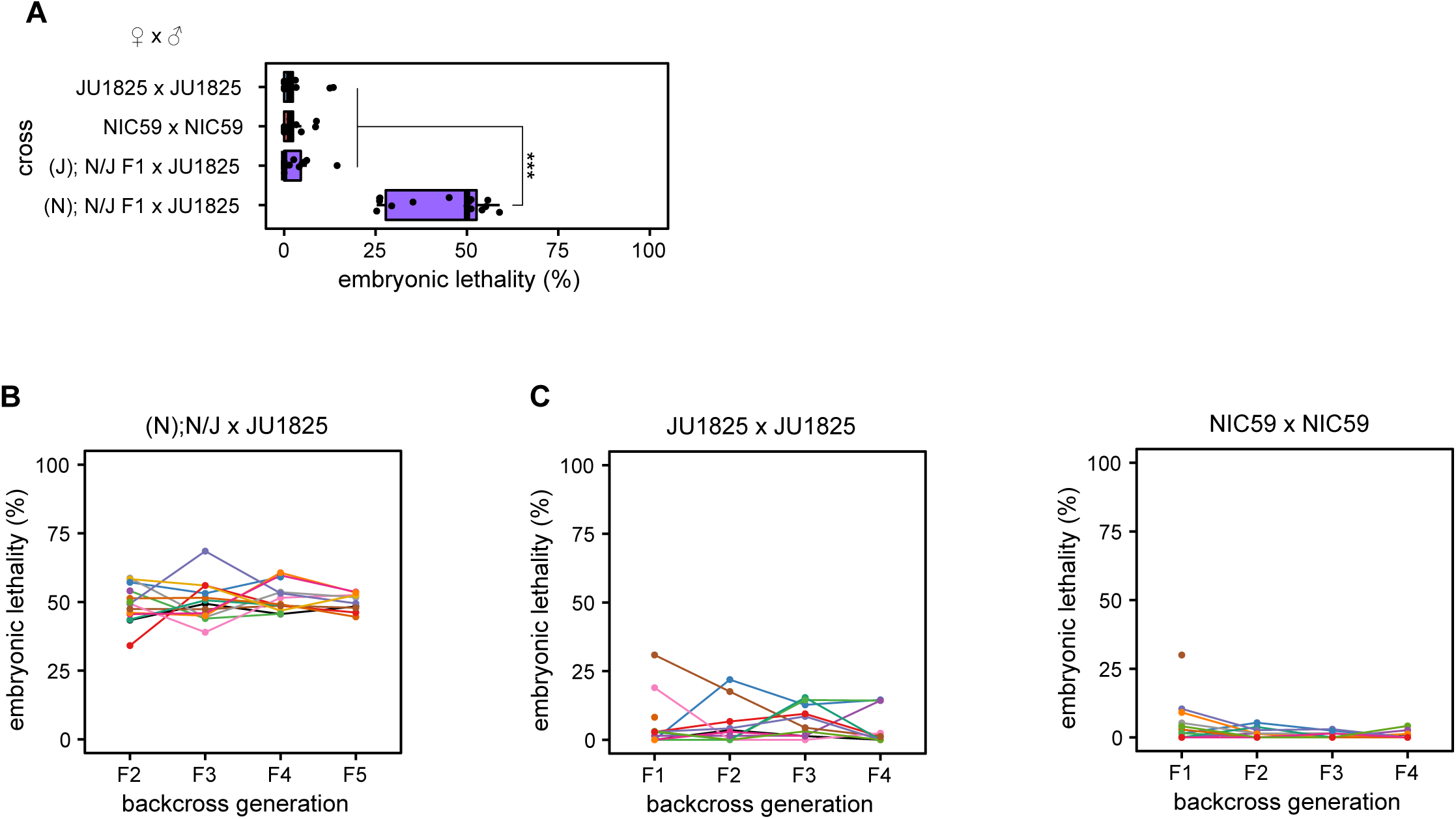
A single BDM incompatibility between a NIC59 cytoplasmic locus and a JU1825 nuclear locus causes embryonic lethality. **(A)** Approximately 50% of the F2 progeny from (N); N/J F1 female x JU1825 male crosses arrest during embryogenesis, significantly higher than that seen in intra-strain crosses (P<0.001). In contrast, (J); N/J F1 female x JU1825 male and parental strain crosses exhibit low and similar levels of embryonic lethality (P>0.05). **(B)** Initially, fifteen (N); N/J F1 females were independently backcrossed to single JU1825 males. For each independent lineage, a single surviving F2 female was again backcrossed to a JU1825 male. This backcrossing scheme was repeated until the F5 generation (11 lineages remain). Each colored line represents a single backcross lineage. All backcross lineages exhibit ~50% embryonic lethality throughout the backcross generations, consistent with the hypothesis that an incompatibility between a NIC59 cytoplasmic locus and a single JU1825 nuclear locus causes embryonic lethality. **(C)** Both parental strains were "backcrossed” as a negative control (JU1825 =14 backcross lineages in the F1 generation, 11 lineages by the F4 generation; NIC59 = 15 backcross lineages in the F1 generation, 11 lineages by the F4 generation). All p-values were calculated by a Kruskal-Wallis test followed by Dunn’s test.

To test the hypothesis of a single BDM incompatibility, we crossed viable F2 females to JU1825 males and assayed F3 viability. Under this hypothesis, the surviving F2 females are expected to have inherited NIC59 mtDNA and be heterozygous (i.e. JU1825/NIC59) at the JU1825 nuclear incompatibility locus (Figure 6A). Therefore, crossing these F2 females to JU1825 males should also result in ~50% embryonic lethality in the F3 generation. This pattern should also be true for additional backcross generations (e.g. F4, F5 etc.). Thus, we generated 15 independent backcross lineages, each consisting of matings between single surviving hybrid females and JU1825 males, and monitored each lineage’s viability for four backcross generations. Indeed, the approximately 50% embryonic lethality observed in the F2 generation is also observed in the subsequent backcross generations in all lineages (Figure 4B). These results are consistent with the hypothesis that embryonic lethality is the result of a simple two-locus BDM incompatibility between a purely maternally inherited cytoplasmic NIC59 locus and a single nuclear locus homozygous for JU1825 alleles. We hypothesize that the post-embryonic inviability may be a genetically separable phenotype.

### The JU1825 cytoplasm appears to be heteroplasmic

As previously discussed, the backcross that combines the JU1825 cytoplasmic genotype with the NIC59 nuclear genotype (i.e. (J); N/J F1 female x NIC59 male crosses) results in ~50% F2 viability on average (Figure 2B). Thus, the total F2 inviability could be the result of a single BDM incompatibility between a JU1825 cytoplasmic locus and a single autosomal locus homozygous for NIC59 alleles. To test this hypothesis, we generated 15 independent backcross lineages, each consisting of matings between single surviving hybrid females and NIC59 males, and monitored each lineage’s viability for four backcross generations. To our surprise, while some lineages continued to exhibit low levels of viability similar to the F2 generation average (~50%), others began to exhibit and maintain significantly increased viability for multiple backcross generations (Figure 5A). For example, in this particular experiment we found that in the F2 generation a majority of lineages (13/15) had a total viability ranging from 18-50%, while only two exhibited higher viability (68% and 85%). However, by the F5 backcross generation, we found that of the fourteen remaining lineages only four exhibited 50% viability or less. Strikingly, by the F5 generation, 5/14 backcross lineages exhibited nearly 100% viability.

**Figure 5.**
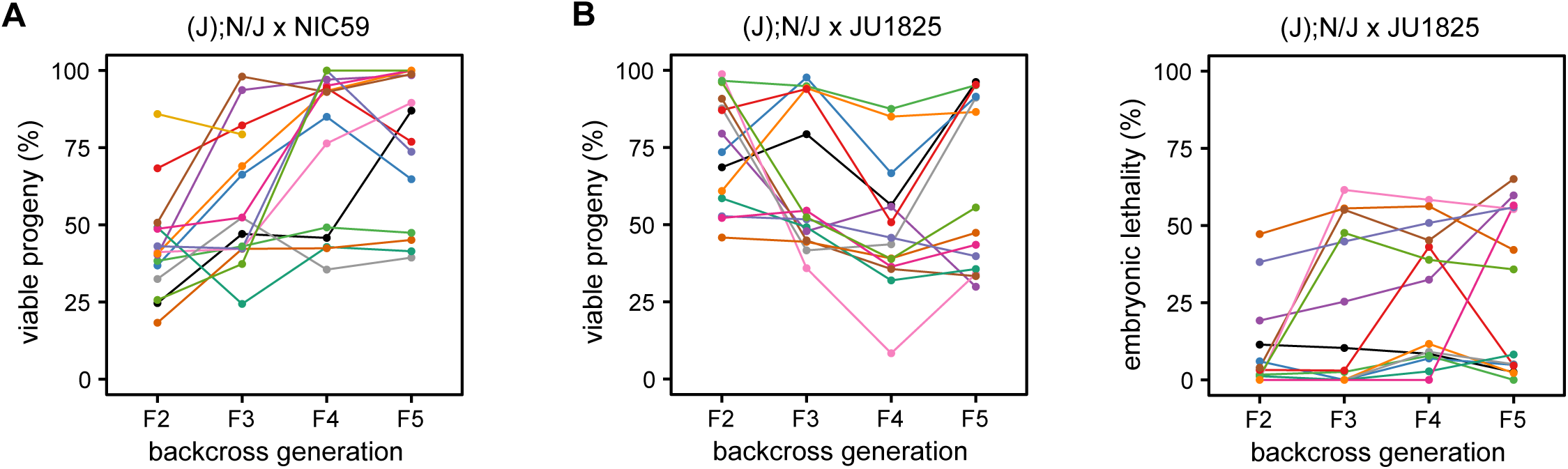
The JU1825 cytoplasm is heteroplasmic for JU1825-like and NIC59-like alleles. **(A)** The viability of 15 independent (J); N/J female x NIC59 male backcross lineages were followed until the F5 generation. Surprisingly, in some lineages, multiple generations of backcrossing resulted in increased viability (similar to that seen in intra-strain crosses). **(B)** The viability of 15 independent (J); N/J female x JU1825 male backcross lineages were also followed until the F5 generation. Interestingly, multiple generations of backcrossing results in some lineages with significantly reduced viability, similar to that seen in (N); N/J F1 female x JU1825 male crosses. We also examined embryonic lethality, finding that some lineages exhibit the ~50% embryonic lethality seen in (N); N/J F1 female x JU1825 male crosses upon additional generations of backcrossing. These results are consistent with the hypothesis that the JU1825 cytoplasm is heteroplasmic and contains JU1825-like and NIC59-like alleles.

The rescue of hybrid inviability for some lineages via several generations of backcrossing is peculiar. One hypothesis to explain this phenomenon is that the JU1825 cytoplasmic and/or NIC59 nuclear incompatibility loci are not fixed within their respective strains, but rather are segregating polymorphisms. As a specific example, the JU1825 cytoplasmic incompatibility locus could be heteroplasmic for alleles that are either incompatible or compatible with the NIC59 nuclear genome. The mitochondrial genome is present at a high copy number within a single cell, and it is thought that individual mtDNAs are randomly replicated and segregated to daughter cells during cell division. Studies on the inheritance of various mtDNA heteroplasmies show that their frequency amongst siblings from the same mother can be highly variable due to the random sampling of mtDNAs and genetic bottlenecks during female germline development (Wallace and Chalkia 2013). Therefore, it is possible that a NIC59-compatible cytoplasmic allele has increased in frequency in some backcross lineages and rescued inviability.

To gain a better understanding of the genetic composition of the JU1825 cytoplasm, we also monitored the viability of (J); N/J female x JU1825 male lineages over four backcross generations. Because this cross combines homotypic JU1825 cytoplasmic and JU1825 nuclear genotypes, we originally predicted that the relatively high rates of F2 viability would persist or possibly increase with additional backcross generations. However, we instead observed that some backcross lineages showed a striking decrease in viability after the F2 generation (Figure 5B). For example, in this particular experiment, lineages in the F2 generation exhibited a uniform distribution of viability, with an average of 74%. By the F5 generation we find two distinct populations of lineages, those with a high viability ranging from 85-96% (6/14 lineages) and those with an astonishingly low viability ranging from 29-55% (8/14 lineages) (Figure 5B). The latter population has an average viability of 39%, which is quite similar to that observed in (N); N/J F1 female x JU1825 male crosses (~36%, Figure 2B), indicating that although these lineages inherited their cytoplasm from JU1825 mothers, they now seem to exhibit low levels of viability similar to those observed in the NIC59 cytoplasmic – JU1825 nuclear incompatibility. One hypothesis to explain these data is that the JU1825 cytoplasm harbors a NIC59-like allele which at a certain threshold frequency can mimic the NIC59 cytoplasmic-JU1825 nuclear incompatibility in certain (J); N/J F1 female x JU1825 male backcross lineages.

In support of this hypothesis, the rate of embryonic lethality for some (J); N/J female x JU1825 male backcross lineages also increases to levels observed in the NIC59 cytoplasmic – JU1825 nuclear incompatibility (i.e. 50%) and can be stably inherited for several backcross generations (Figure 5C). Specifically, most lineages (12/14) in the F2 generation exhibited only 0-19% embryonic lethality, while only two lineages exhibited higher rates (38 and 47%). However, by the F5 backcross generation, only about half of the lineages (6/14) exhibited 0-8% embryonic lethality, while 8/14 lineages exhibited 35-65% embryonic lethality. Taken together, the results from the two backcross experiments are consistent with the hypothesis that the JU1825 cytoplasm is heteroplasmic and harbors both JU1825-like and NIC59-like incompatibility loci (Figure 6B and C).

**Figure 6.**
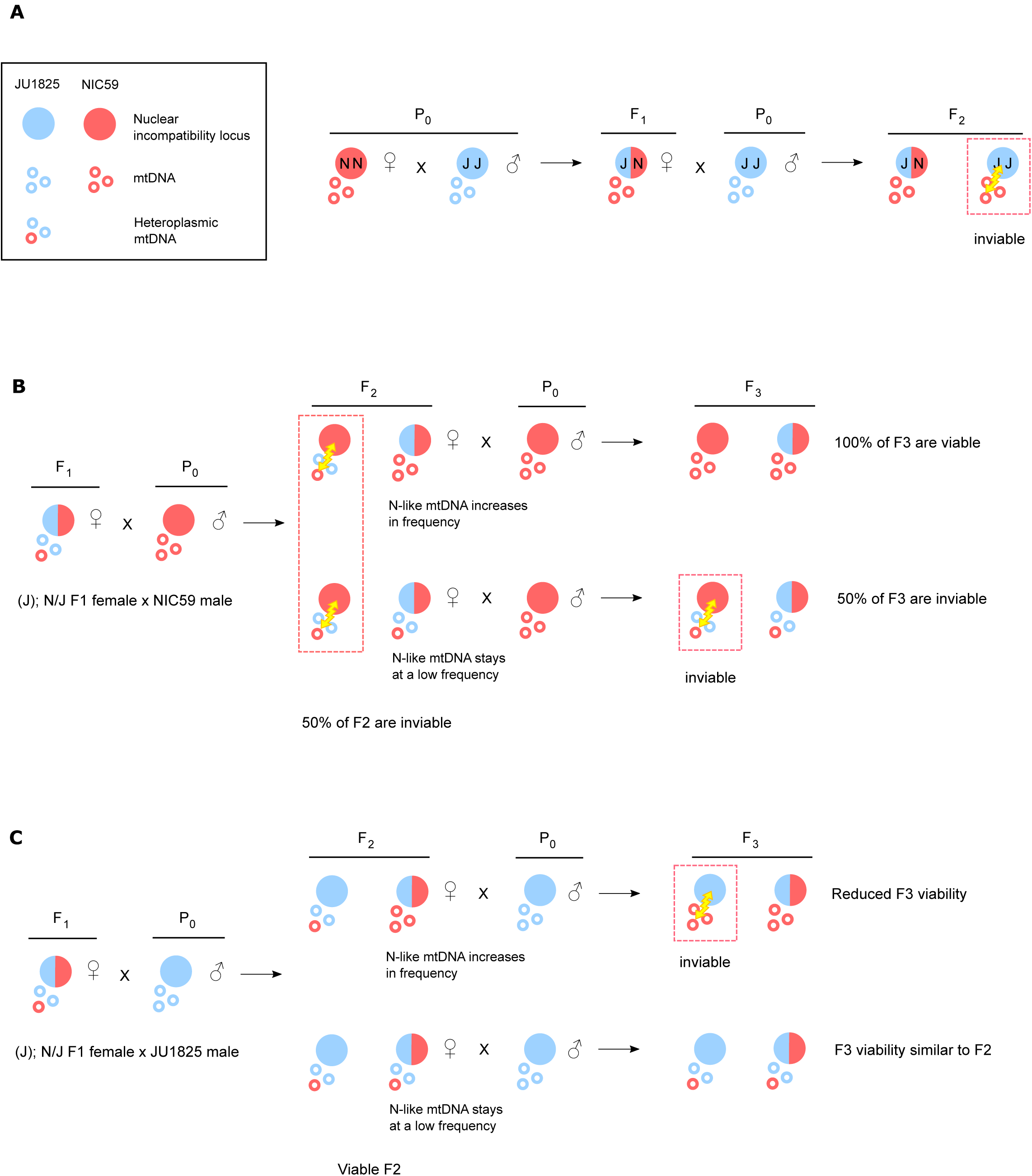
Mitochondrial-nuclear incompatibility model. **(A)** We hypothesize that F2 hybrid breakdown is the result of a Bateson-Dobzhansky-Muller incompatibility between the NIC59 mitochondrial genome and a nuclear locus homozygous for the JU1825 allele, and vice versa. As a specific example, when NIC59 females are crossed to JU1825 males, the resulting F1 hybrid females are expected to be heterozygous at all autosomal loci while inheriting only NIC59 mtDNA. When F1 females are backcrossed to JU1825 males, F2 inviability results from an incompatibility between NIC59 mtDNA and an autosomal locus homozygous for the JU1825 nuclear allele. **(B)** We hypothesize that the JU1825 cytoplasm is heteroplasmic in F1 females and contains at least one NIC59-like allele. Backcrossing hybrid females with a JU1825 cytoplasm (i.e. (J); N/J females) to NIC59 males for multiple generations can allow the NIC59-like cytoplasmic allele to increase in frequency and dilute out the effects of the incompatible JU1825-like mtDNA (e.g. top right F2 female). This eventually may allow the NIC59 nuclear locus to become homozygous and restore the viability of a lineage. On the other hand, the NIC59-like mtDNA can stay at a low frequency in viable F2 females (e.g. bottom right F2 female). Backcrossing these F2 females to NIC59 males results in levels of inviability similar to the F2 generation. **(C)** By a similar line of reasoning, backcrossing hybrid females with a JU1825 cytoplasm to JU1825 males for multiple generations can allow the NIC59-like mtDNA to increase in frequency, where it can mimic the same genetic incompatibility seen in (N); N/J F1 female x JU1825 male crosses (Figure 6A).

## DISCUSSION

We discovered a lethal cytoplasmic-nuclear incompatibility between two wild isolates of *C. nouraguensis*, JU1825 and NIC59, and find that such incompatibilities may be widespread between other wild-isolates within the species. We show that maternally inherited endosymbiotic bacteria are not the cause of hybrid inviability, making the mitochondrial genome the most likely candidate for harboring the cytoplasmic incompatibility factor(s). We also propose that the JU1825 cytoplasm is heteroplasmic, and harbors both JU1825-like and NIC59-like incompatibility loci.

In eukaryotes, the mitochondrial genome typically contains a very small fraction of the gene content of a cell, yet it seems to be involved in a disproportionate number of genetic incompatibilities across a diverse range of taxa (Rand *et al.* 2004; Burton and Barreto 2012). However, there are relatively few cases in which incompatibility loci have been definitively mapped to the mitochondrial genome, and therefore a larger sample is required to better understand what drives the evolution of mitochondrial-nuclear incompatibility. Additionally, all of the molecularly identified cases of mitochondrial-nuclear incompatibility have been found between species rather than within species (Lee *et al.* 2008; Chou *et al.* 2010; Luo *et al.* 2013; Meiklejohn *et al.* 2013; Ma *et al.* 2016). Some of these inter-species hybridizations harbor additional genetic incompatibilities or chromosomal rearrangements that cause inviability and sterility (Hunter *et al.* 1996; Fischer *et al.* 2000; Brideau *et al.* 2006; Ferree and Barbash 2009; Mihola *et al.* 2009; Davies *et al.* 2016), making it difficult to discern whether mitochondrial-nuclear incompatibility was instrumental in initiating speciation or evolved after strong reproductive isolation occurred. The incompatibility we describe here provides an excellent opportunity to study the evolutionary genetics and cell biology of incipient speciation as well as mitochondrial-nuclear incompatibility. The ease of breeding, large brood sizes, and short generation time of *C. nouraguensis* should facilitate the mapping and identification of the genes that contribute to hybrid inviability.

### Asymmetric cytoplasmic-nuclear incompatibilities

Most molecularly characterized BDM incompatibilities are asymmetric, in that only one of two divergent alleles at a locus is incompatible with heterospecific alleles at other loci (e.g. Brideau *et al.* 2006; Ferree and Barbash 2009). This is also true of the mitochondrial-nuclear incompatibilities seen in *Saccharomyces* species hybridizations (Lee *et al.* 2008; Chou *et al.* 2010). For example, an intron of the *COX-1* gene in the *Saccharomyces bayanus* mitochondrial genome fails to be correctly spliced by the nuclearly encoded S. *cerevisiae MRS-1* gene, resulting in hybrid inviability presumably due to a failure of respiration on non-fermentable media. However, a similar incompatibility does not occur between S. *cerevisiae COX-1* and S. *bayanus MRS-1.* At first glance it may seem as though the cytoplasmic-nuclear incompatibility between JU1825 and NIC59 is symmetric, since both strains have a cytoplasm that is incompatible with the other strain’s nuclear genome. However we currently cannot determine whether the JU1825 or NIC59 cytoplasmic and nuclear incompatibility loci are allelic, leaving open the possibility that different genes cause hybrid inviability in the reciprocal crosses.

### Cytoplasmic-nuclear incompatibility: both sexes are equally inviable

J.B.S Haldane noted that the heterogametic sex more often suffers from inviability or sterility in inter-species hybridizations than the homogametic sex (Delph and Demuth 2016). It is not known whether “Haldane’s rule” also applies in intra-species hybridizations. The lethal cytoplasmic-nuclear incompatibility we identified between the NIC59 and JU1825 wild isolates of *C. nouraguensis* affects females and males equally, suggesting that the two sexes share the same disrupted developmental pathway(s). However, we have not carefully studied other aspects of sex-specific fitness, such as female and male F2 hybrid fertility. Because the mitochondrial genome is inherited only through females, theory predicts that evolution will lead to the accumulation of mtDNA variants that are neutral or increase female fitness, but that are neutral or possibly deleterious to male fitness (Gemmell *et al.* 2004). Because of this, male-specific functions may be more adversely affected during the hybridization of heterotypic mitochondrial and nuclear genomes. This is indeed the case for some known mitochondrial-nuclear incompatibilities. For example, when swapping the mitochondrial genomes between mouse subspecies via pronuclear transfer, one mitochondrial-nuclear combination resulted in reduced male fertility while females had relatively normal fertility (Ma *et al.* 2016). Therefore, further studies of *C. nouraguensis* hybrid male fertility are required to more fully address whether this system follows Haldane’s rule, as well as to determine whether there are male-specific mitochondrial-nuclear incompatibilities.

### JU1825 heteroplasmy

We hypothesize that the JU1825 cytoplasm is heteroplasmic and contains mitochondrial genomes that are both compatible (JU1825-like) and incompatible (NIC59-like) with the JU1825 nuclear incompatibility locus. If the JU1825 cytoplasm is naturally heteroplasmic, we predict the NIC59-like mtDNAs are kept at a low frequency within JU1825 by selection. This selection would be relaxed in (J); N/J F1 hybrids and the frequency of NIC59-like mtDNA can increase, reducing incompatibility in backcrosses to NIC59 males and increasing incompatibility in backcrosses to JU1825 males. However, our data cannot rule out the possibility that NIC59-like mtDNA is introduced into F1 females by incomplete degradation and inheritance of paternal NIC59 mtDNA. Interestingly, indirect evidence suggests that paternal mtDNA can be inherited when hybridizing different wild isolates of *Caenorhabditis briggsae* (Hicks *et al.* 2012; Chang *et al.* 2015).

The hypothesized heteroplasmy of the JU1825 cytoplasm may explain the greater variance of F2 viability in crosses with (J); N/J F1 females in comparison to those with presumably homoplasmic (N); N/J F1 females. Stochastic segregation and genetic bottlenecking events from JU1825 mothers may result in F1 females with a wide range of frequencies of the NIC59-like cytoplasmic allele, and therefore a wide range of F2 viability when backcrossed to either NIC59 or JU1825 males. This stochastic inheritance may also explain why the degree of F2 viability of (J); N/J F1 female x JU1825 male backcrosses can also vary significantly from experiment to experiment (Figure S1).

## Acknowledgments

We thank Marie-Anne Félix and Christian Braendle for providing the *Caenorhabditis nouraguensis* strains used in this study. We also thank the labs of Harmit Malik and Leo Pallanck for providing *Drosophila melanogaster* strains with and without *Wolbachia.* We thank Janet Young, Harmit Malik, Maitreya Dunham and Irini Topalidou for helpful discussions and comments on the manuscript. P.L. was supported in part by an NIH Institutional Training Grant (PHS, NRSA, T32GM007270 from NIGMS). This work was supported by an NSF CAREER award to M.A.

## Figure Legends

**Supplemental Figure 1. Variability of (J); N/J F1 female x JU1825 male crosses across experiments.** Three biological replicates of the same type of backcross experiment. (J); N/J F1 female x JU1825 male crosses can either exhibit similar or significantly decreased rates of viability in comparison to intra-strain crosses across experiments (Experiment 1, non-significant, P>0.05; Experiment 2, non-significant, P>0.05; Experiment 3, P>0.05, non-significant in comparison to JU1825 x JU1825 crosses, P<0.05 significant in comparison to NIC59 x NIC59 crosses). However, (J); N/J F1 female x JU1825 male crosses consistently exhibit significantly increased rates of viability in comparison to (N); N/J F1 female x JU1825 male crosses (Experiment 1, **, P<0.01; Experiment 2, **, P<0.01; Experiment 3, *, P<0.05). Experiments 1 and 2 are data from Figure. 2 and 5, respectively. All p-values were calculated by a Kruskal-Wallis test followed by Dunn’s test.

**Supplemental Figure 2. Nuclear incompatibility loci are linked to autosomes, not sex chromosomes.** F1 intercrosses allow us to infer that the nuclear incompatibility loci are autosomal, not X-linked. From the (N); N/J F1 x JU1825 male backcross experiment (Figure 2), we concluded that F2 inviability was the result of a genetic incompatibility between the NIC59 mitochondrial genome and nuclear loci homozygous or hemizygous for JU1825 alleles. It is reasonable to assume that the same genetic incompatibility contributes to F2 inviability in (N); N/J F1 female x (N); N/J F1 male crosses. If the nuclear incompatibility locus were X-linked, F2 males would have a 50% chance of being hemizygous for the JU1825 nuclear incompatibility locus while F2 females can only be heterozygous or homozygous for NIC59 alleles. Therefore, if the locus were X-linked, half of F2 males would be inviable while females would be unaffected. If the nuclear incompatibility locus were autosomally linked, then both sexes have an equal chance of being homozygous for the JU1825 nuclear incompatibility locus. Therefore, both sexes are expected to suffer equal rates of inviability. We do not observe a significant decrease in the proportion of viable F2 males (Figure 1), so we conclude that the JU1825 nuclear incompatibility locus or loci are linked to autosomes. The same line of reasoning can be used to show that the NIC59 incompatibility locus or loci are also autosomally linked.

**Supplemental Figure 3. Endosymbiotic bacteria do not cause cytoplasmic-nuclear incompatibility. (A)** PCR on both JU1825 and NIC59 crude lysates (10 adult worms per lysate, 5 females and 5 males) with degenerate primers against the *Wolbachia fbpA* or *gatB* loci fails to amplify the expected products. w^1118^ (wol+) and w^1118^ (wol-) *D. melanogaster* flies serve as positive and negative controls, respectively. PCR on crude lysates of OP50 (bacterial food source of NIC59 and JU1825) also fails to amplify the expected products. PCR on JU2079, an inbred strain derived from JU1825, also fails to amplify the expected *gatB* product. **(B)** Both tetracycline-treated (J);N/J F1 female x NIC59 male and (N);N/J F1 female x JU1825 male crosses exhibit significantly decreased levels of viability in comparison to tetracycline treated intra-strain crosses (P<0.01). Additionally, there are no statistical differences in viability between NIC59 x NIC59 and JU1825 x JU1825 tetracycline treated intra-strain crosses (P>0.05). N=14 or 15 for each cross. All p-values were calculated by a Kruskal-Wallis test followed by Dunn’s test.

**Supplemental Figure 4. Cytoplasmic-nuclear tests separated by nuclear genotype.** Each graph depicts all the cytoplasmic-nuclear tests performed between four cytoplasmic genotypes and a single nuclear genotype. This is the same data that is grouped into a single graph in Figure 3B. Each cytoplasmic-nuclear combination has three biological replicates (except for JU1825 cytoplasm – NIC24 nuclear and JU1825 cytoplasm – NIC54 nuclear combinations, which have four replicates). Although there appear to be many cases of significant cytoplasmic-nuclear incompatibility (relative viability<1 and P<0.0006 after Bonferroni correction), there can be discrepancies between replicates (e.g. one replicate of the JU1825 cytoplasm - NIC24 nuclear combination indicates a significant incompatibility, while the other three do not).

